# Calculation of minimum energy pathways in transport proteins

**DOI:** 10.1101/2024.08.07.607056

**Authors:** Briony A Yorke, Helen M Ginn

**Author notes:** Corresponding author’s.

## Abstract

Diverse conformations of highly populated protein metastable states are well-studied, but the fleeting transitions between these states cannot be observed by experimental methods or molecular dynamics simulations. To address this, we present a computationally inexpensive algorithm, “cold-inbetweening”, which generates trajectories in torsion angle space, by minimising the overall kinetic energy needed to complete a transition between experimentally determined end-states. Here we demonstrate the application of cold-inbetweening to provide mechanistic insight into the ubiquitous alternate access model of operation in three membrane transporter superfamilies. The model proposes mutually exclusive outward and inward pore opening, allowing ligand translocation but preventing damage from free solvent flow. Here, we study DraNramp from *Deinococcus radiodurians*, MalT from *Bacillus cereus*, and MATE from *Pyrococcus furiosus*. In MalT, the trajectory demonstrates elevator transport through unwinding of a supporter arm helix, maintaining adequate space to transport maltose. In DraNramp, outward-gate closure occurs prior to inward-gate opening, in agreement with the alternate access model. In the MATE transporter, switching conformation involves obligatory rewinding of an extended N-terminal helix to avoid steric backbone clashes. This concurrently plugs the cavernous ligand-binding site during the conformational change. More generally, cold-inbetweening can be used to inform hypotheses about large functionally relevant conformational changes.

## 2 Introduction

Neither molecular dynamics simulations nor experimental methods provide sufficient information about large transitions between conformational states in proteins to fully characterise their mechanisms. The reason for this is that events occur quickly but infrequently and at random times, making them very hard to study. The transition time can be orders of magnitude smaller than the time taken waiting for this to occur. There is a need for computational calculation of trajectories, for which previous efforts [1–5] have made incremental progress in this regard. Others are considered aesthetical rather than analytical tools [6]. Each of these methods require the researcher to supply two trusted structures to serve as end-points, and from these, interpolated structures are hypothesised. Due to limitations of existing methods, the way to and from these end-points remain unexploited. To address this, we have developed an algorithm, cold-inbetweening, to generate trajectories that smoothly connect starting and ending conformations. The algorithm mimics the nature of protein flexibility by allowing rotation around bonds, since the energy needed to change bond lengths, and to a lesser degree bond angles, is far larger than that needed to change torsion angles. Additionally, the algorithm is designed to minimise fluctuations in kinetic and potential energy needed to complete a transition between experimentally determined end-states, thereby adhering to the third of Newton’s laws of motion. The model of motion omits random thermal fluctuations other than those associated with the desired conformational change.

To test the cold-inbetweening algorithm we applied it to the analysis of transport protein mechanisms from three distinct superfamilies to investigate whether the calculated trajectories were in agreement with the literature.

Transport proteins are able to move specific molecules across a membrane against a concentration gradient. In general, they are thought to function according to the hypothetical alternate access mechanism [7, 8]. According to the alternate access model, the protein binding site is exposed at one side of a membrane and can non-covalently bind a ligand. The binding event then initiates a conformational change that translocates the ligand through the membrane. The binding site is then subsequently exposed to the other side of the membrane and the ligand released. The alternate access mechanism is further sub-categorised into the elevator and rocking-bundle mechanisms [9]. The elevator mechanism refers to the movement of domains after ligand-binding such that the domain-domain interface changes substantially and the protein-ligand interface is largely unchanged [10]. In contrast, the rocking bundle mechanism involves a domain motion that substantially changes the protein-ligand interface itself. The binding site is enclosed between two rigid bodies, and is the fulcrum around which one of the rigid bodies rotates, thereby alternating access between the two sides of the membrane [11].

Transport proteins are particularly suited to our method since they often have a few dominant conformations, inward-open (where the binding site is exposed to the inner side of the membrane),

outward-open (where the binding site is exposed to the outer side of the membrane) and occluded (where the binding site is shielded from both sides of the membrane). However, the details of the motion involved in these mechanisms are unknown. We selected three classes of conformational change to test the cold-inbetweening algorithm. Specific transport proteins were chosen according to the availability of high-quality models of the inward-open and outward-open states in the protein databank to use as our starting and ending conformations.

## 3 Results and Discussion

The algorithm assumes regularised bond lengths, angles and optimised torsion angles [12] of both starting and ending structures. Close pairs of atoms in the protein structure are considered. Each pair has a starting and ending inter-atomic distance. A glidepath for these distances during the trajectory is sought that minimises the fluctuations in kinetic and potential energy of the whole protein during the transition (Fig. 1A). To minimise these fluctuations (such as Van der Waals and electrostatic energy terms), the ideal glidepath would be linear. However, this is not fully achievable due to the constraints imposed by limited freedom of motion in torsion angle space. Therefore, the torsion angle motions are adjusted to minimise the differences between the actual interatomic distances and their ideal ones (Fig. 1B). Initially, torsion angles take a linear course between values from the start and end structure. Harmonic perturbations are added with refinable amplitudes to minimise deviation from the ideal interatomic paths (called early-stage refinement, see Methods). Residual clashes in sidechains are then resolved by minimising a combined energy term (called late-stage refinement, see Methods).

**Fig. 1:**
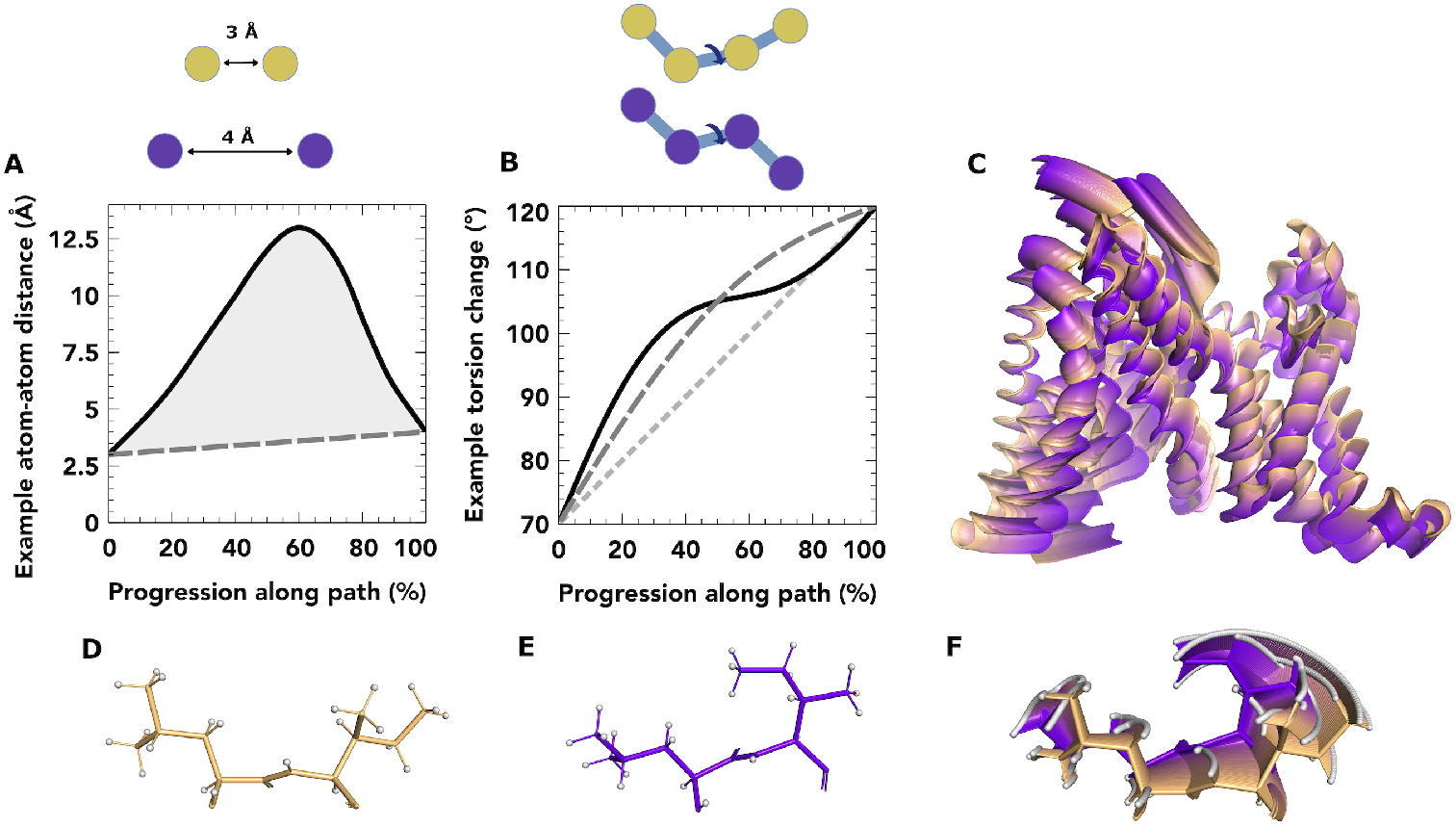
(A) illustrative atom-atom distance term of momentum target function showing torsion angle-mediated distance (black solid), and target glidepath (dark grey dashed). Minimisation reduces shaded area. (B) torsion angle trajectories are parameters, starting from linear interpolation (light grey dotted), addition of a first-order harmonic (dark grey dashed) and second-order (black solid). Relative contributions of perturbations serve as parameters. (C) after refinement of MalT transporter path, overall motion rendering the entire pathway from outward-open (orange) to inward-open (purple) in 1% steps. (D-F) zoom-in of residues 385 and 386, showing (D) start-, (E) end-state and (F) all 1% steps superimposed. (C-F) figures generated in PyMOL [26].

### 3.1 Elevator mechanism of maltose transport

The EIIC domain of MalT mediates the transport of maltose and is purported to act by the elevator subcategory of alternate access. The structure of this protein has been solved in an outward-open (PDB: 6bvg) and inward-open (PBD:5iws) conformation [13]. This was the simplest rearrangement in our study and we use this to demonstrate some characteristics of the method. We show the cold-inbetweened path in Fig. 1C. Use of harmonic adjustments allows backbone atoms to follow near-perfect straight lines, despite moving by rotation around bonds. Sidechains, often undergoing much larger motions than the backbone, often take more curvaceous routes due to restrictions of that movement around a limited number of bonds (Fig. 1D-F).

The full path shows the coupled winding of helix AH2 to become an extension of TM5, and a subsequent collapse of the angle between AH2 and TM6 (Supporting Movie 1). Despite the method not explicitly modelling a ligand, we found that it is nevertheless suitable to maintaining the binding-cavity (Fig. 2A). This is because the starting and ending pairwise distances between atoms encode the size of the ligand-binding cavity, and there is no term in the target function which would lead to its collapse. Two residues V20 and V353 also appear to act as a gate, closing the maltose molecule off from the solvent during transit (Fig. 2B,C). The cavity is also maintained by miniature unwinding events in helix TM1, which forms one of the three walls of the elevator (Supporting Movie 1).

**Fig. 2:**
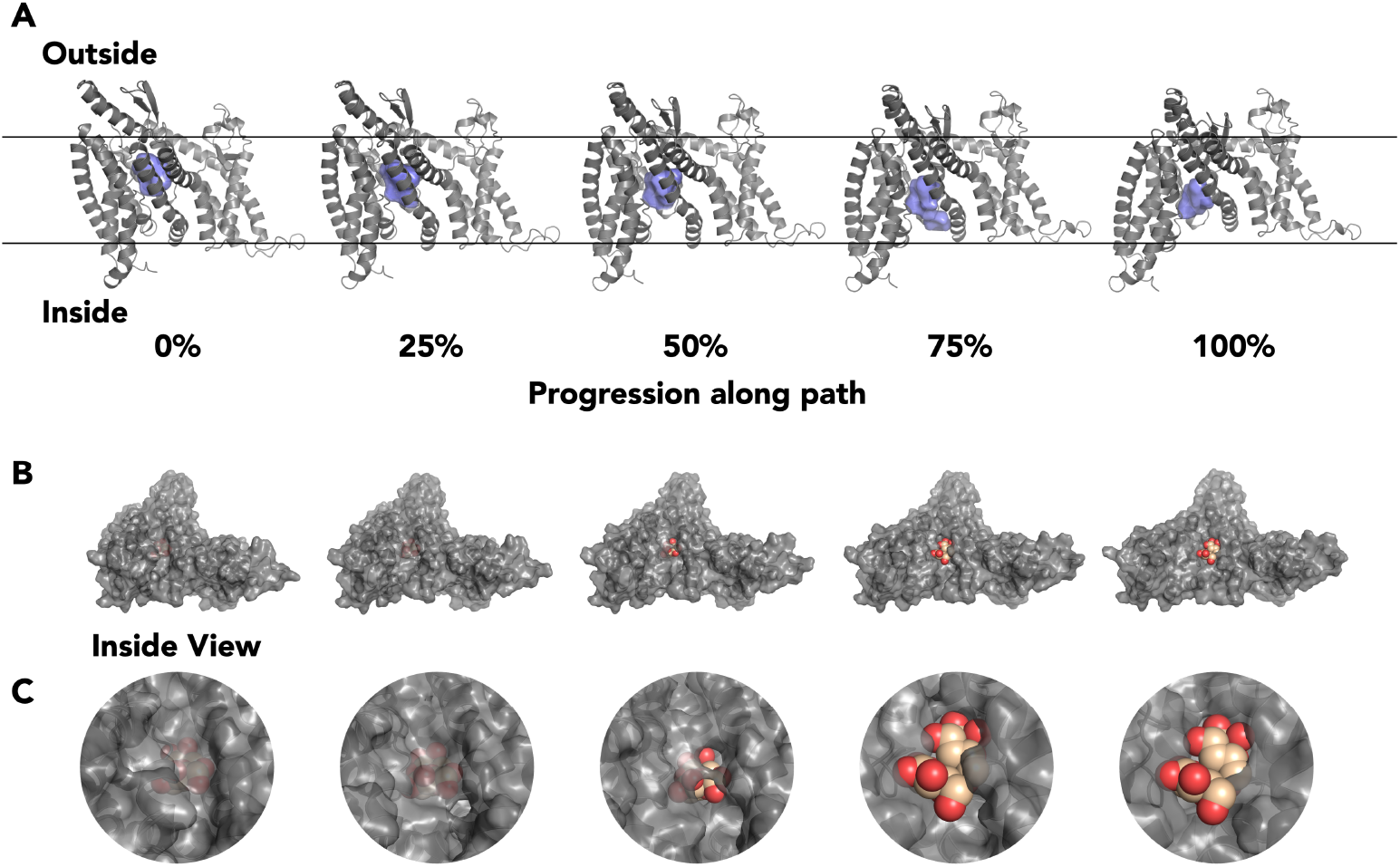
(A) movements of the binding cavity in the EIIC domain of the MalT transporter, in 25% intervals along the trajectory with 0% and 100% corresponding to outward-open and inward-open respectively. In each figure the binding cavity, shown as a blue surface, was calculated using KVfinder [27]. (B) inside view of MalT showing movement of maltose (orange) in the binding cavity and opening of the inward pore. At 0% progression the maltose co-ordinates were taken directly from the outward open stucture (PDB 5iws), for progressive frames the maltose was manually translated to fit inside the cavities using PyMOL [26]. (C) Close-up of the inside pore opening of MalT and movement of maltose.

### 3.2 Rocking bundle mechanism of a LeuT fold protein

Next we applied the algorithm to less straightforward rearrangements. NRAMP proteins transport metal ions in a wide array of organisms. In humans, mutations of NRAMP proteins lead to immune deficiency and anaemia. DraNramp is a tractable bacterial homologue of human NRAMP from *Deinococcus radiodurians* and a member of the ubiquitous LeuT superfamily [14, 15]. DraNramp transports manganese and other heavy metal ions from outward-to inward-facing sides of the membrane. The first structure of DraNramp naturally crystallised in the inward-open form [16]. By introducing steric bulk in the outward-gate (G223W), DraNramp was crystallised in an outward-open form [17]. This mutation also impaired metal transport and prevented NEM modification of an inward-gate reporter site (A53C), demonstrating inward-gate closure does not occur if the outward gate is forced open [17]. This therefore supports the alternate access mechanism. LeuT fold proteins are purported to work by the rocking-bundle subcategory of alternate access [18, 19]. However, for all LeuT transporters, direct visualisation of the transition from inward-to outward-facing structures has, as for most large-scale conformational changes, been so far unattainable. Many solved structures of DraNramp have unmodelled loops. As a continuous chain of torsion angles is required for cold-inbetweening, we chose the fully-modelled Mn(II)-bound outward-open and inward-open conformations as start and end-points (PDBs 8e6n and 8e6i respectively) [20]. To preserve the original sequence, A230 was mutated to wildtype methionine manually altered to match the rotamer from the discontinuous apo-inward-open structure (PDB 6d9w) [16]. We omitted the inward-occluded structure as it was unclear to what extent the G45R mutation was distorting the otherwise untethered TM1a helix.

In our pathway, a rapid and consistent contraction of TM10 and TM6a against TM1b closes off access to the ligand-binding site from the outward vestibule (Fig. 3A, Supporting Movie 2). We monitor this using the area between the C*α* atoms of G58, A227 and Q378 (outward triangle), which steadily collapses (Fig. 3B). The inward-facing vestibule is opened by expansion of a gap between TM1a, TM6b and TM8. The inward triangle for the inward-facing vestibule encompasses C*α* atoms of S327, V233 and A53. In Fig. 3A, A53 rapidly moves away from the ligand-binding site through unwinding of the TM1a-b kinked region. However, this produces a shearing motion of TM1a against TM6b and TM8 in the first half of the pathway, which results in no overall change in the area of the outward triangle. It is only at the 50% mark that inward gate-opening picks up pace to allow solvent access to the ligand-binding site (Fig. 3C). This therefore supports previously anticipated behaviour and provides structural details for this hypothesis. The pathway was heavily influenced by constraints on the substantial rearrangement of the TM6b-TM7 linker. Here, I245 could either flip clockwise or anti-clockwise by 180*°* to the other side of the backbone. Q246 must complete a counter turn either 90*°* in one direction or 270*°* in the other (Supporting Movie 2). One of the two residues needs to turn through the interior of the protein while the other must turn towards the solvent, but only Q246 was able to achieve this without substantial, unresolvable clashes on the loop interior. Accommodation of the 270*°* turn led to a substantial mid-trajectory displacement of TM6b. TM6b moves 1.2 Å out of alignment mid-trajectory (Supporting Fig. 1). This suggests the mechanical hypothesis supports previous NEM-labelling data which indicated that TM6b moves substantially during conformational switching [21].

**Fig. 3:**
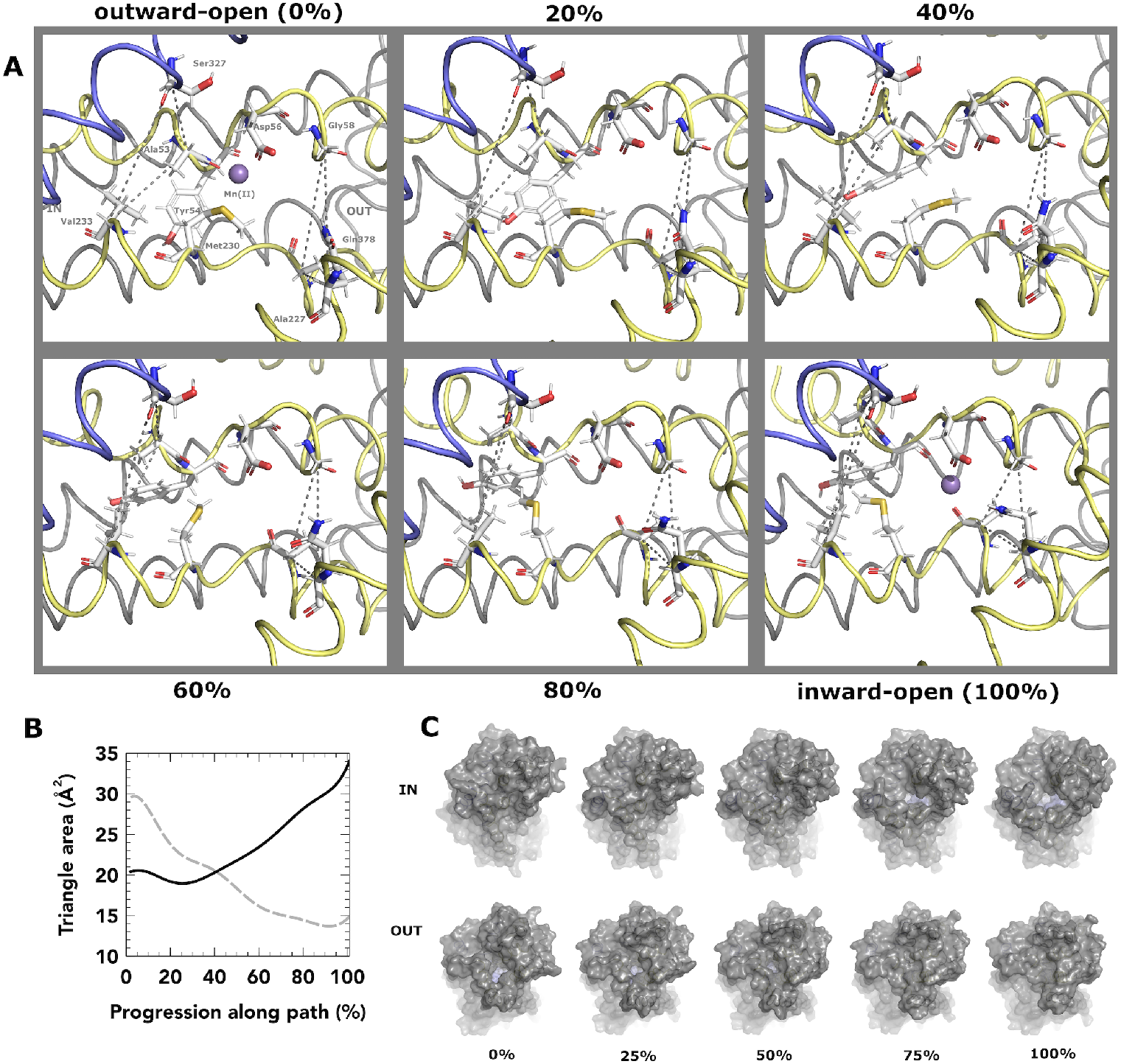
(A) movements at the ligand-binding site (TM helices 3 and 8 removed for clarity) in 20% intervals along the trajectory, with 0% and 100% corresponding to outward-open and inward-open, respectively. In each figure, the left dotted triangle reports on the solvent accessibility to the inward gate, and the right dotted triangle reports on that of the outward gate. (B) area of inward triangle (black solid line) and outward triangle (grey dashed line) along path. (C) surface render from outward and inward sides for five points on path, with residues rendered as sticks in (A) coloured light blue.

### 3.3 Helix unwinding of the MATE transporter

Finally, we examined a member of the MOP superfamily, the multidrug and toxic compound extrusion (MATE) transporter [22]. MATE transports xenobiotics and metabolites across membranes of cells and organelles and in bacteria can confer resistance to antibiotics. MATE from *Pyrococcus furiosus* has a large 4300 Å^3^ cavity. The inward-open and outward-open structures (PDBs 6fhz and 6hfb) [23] differ by a dramatic rewinding of the central portion of the TM1 helix. Firstly, we checked Ramachandran outliers in the unwound structure: a peptide flip of G30 was found to fix a Ramachandran outlier in the inward-open conformation, which fit the density equally well. Other TM1 Ramachandran outliers were favoured in their fit to the density and considered to be realistic in light of the dynamics in this region. The helix, in its unwound state, bridges the large gap between the N-terminal domain and C-terminal domain; whereas the fully-wound helix is able to bridge the narrower gap in the outward-open state. One pertinent question is what combination of bond rotation directions allows the helix to rewind (or unwind) without causing internal steric clashes? By testing several combinations, we found a solution involving a mid-trajectory development and resolution of a kink in the middle of the helix (Fig. 4A, Supporting Movie 3), which did not clash internally or with the rest of the protein (so-called “kinked winding”). This, however, protruded far from the helical axis. We also attempted

**Fig. 4:**
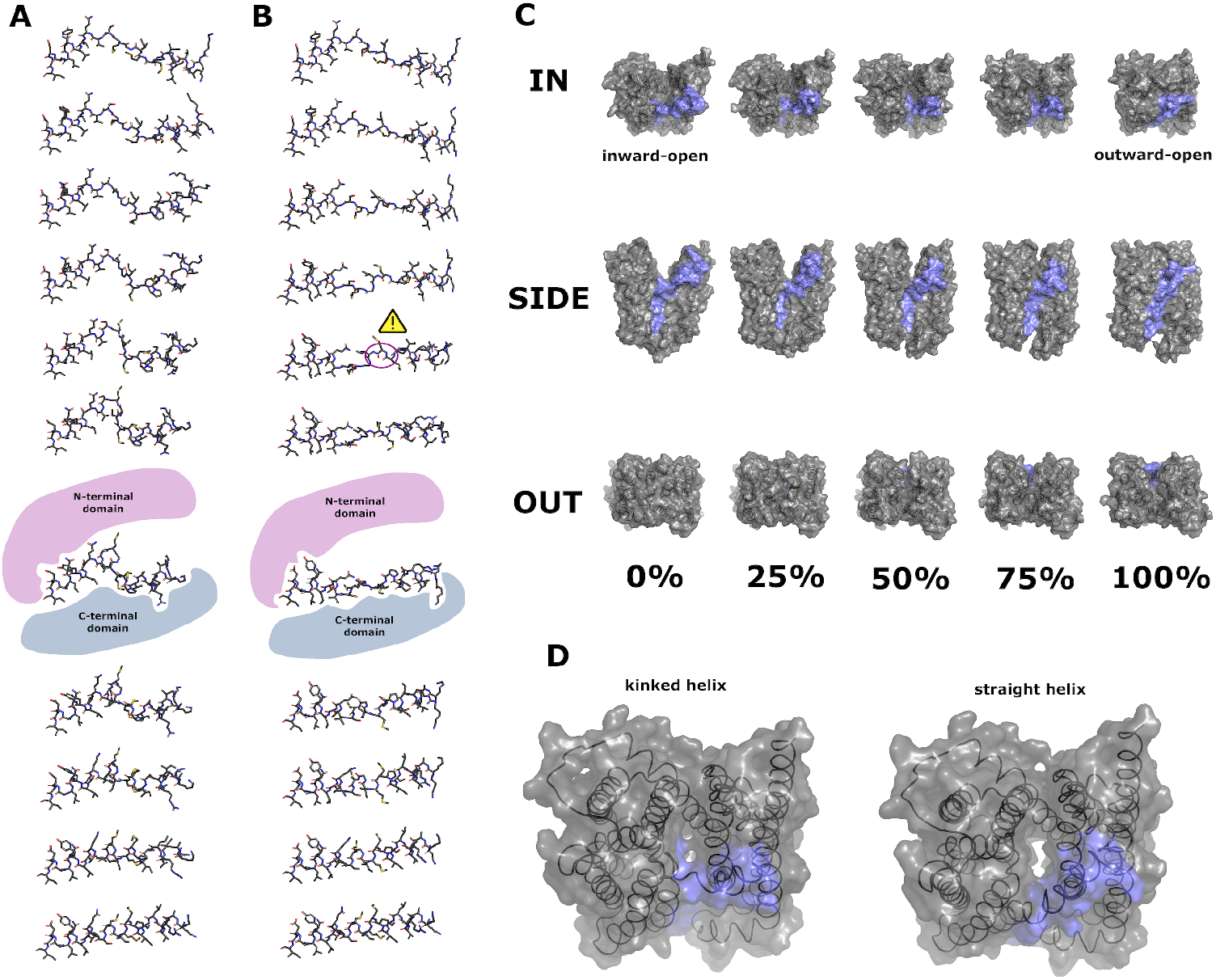
(A) 10% increments in progression from 0% (top) to 100% (bottom) showing atomic trajectory of TM1 with torsion angle flips assigned as necessary to avoid backbone clashes (“kinked helix” trajectory). N-terminus is to the right. (B) similar increments in progression showing atomic trajectory of helix minimising out-of-axis motion (“straight helix” trajectory). Carbonyl clash on backbone highlighted with a warning sign. For (A) and (B) the 60% progression point is augmented by an illustration of relative positions of N-terminal and C-terminal domains. (C) view of kinked helix trajectory from the inward, side and outward views. TM1 residues 1-43 are marked in blue. (D) comparative views of kinked and straight helix trajectories at 70% trajectory, showing the aberrant opening of a solvent channel when all other protein motions with a straight helix trajectory are considered.

a direct recoiling of the extended helix, in order to minimise off-axis motion and thereby conserve momentum (so-called “straight winding”, Fig. 4B). However, we found this would be highly energetically unfavourable due to a close carbonyl-carbonyl clash in adjacent residues (M27 and M28) on the backbone. It also led to a 270*°* rotation of the N-terminal section of TM1 around its axis. However, the kinked winding alternative has a more simple 180*°* shift of this region, which is a more achievable helix register slippage. We also note that at 95% progression through the kinked winding trajectory, the r.m.s.d. of *C*_*α*_ atoms of TM1 reaches a minimum against the bent conformation [24], whereas no mid-trajectory agreement of the straight winding path has a smaller r.m.s.d. than the end-point, strongly suggesting straight winding is not the mechanism by which the alternate access model operates. The kinked winding protrusion extends into the cavity mid-trajectory and thereby blocks solvent access to the cavity (Fig. 4C) which does not occur for straight winding (Fig. 4D). We also checked for overlap of the kinked helix with ligand- and inhibitor-bound structures previously determined [24], and found that overlap with the ligand was insubstantial compared to that of the inhibitors (Supporting Fig. 2).

## 4 Conclusion

Here we show that cold-inbetweening can be used to estimate the feasibility of particular transitions and their consequences. Although thermal motion is the only route by which molecular dynamics (and reality) can escape local energy minima, cold-inbetweening finds a common pathway around which true trajectories are likely to follow, albeit with additional noise attributed to heat. This therefore ought to complement the experimental determination of structures and simulations using the heat-driven engine of molecular dynamics simulations. The omission of thermal motion precludes direct comparison to the results of molecular dynamics simulations, and we do not assign a timescale to the trajectory. We show that for membrane transporter proteins, cold-inbetweening preserves ligand-binding states due to their inclusion in the start and end models even without explicitly modelling an energy term for electrostatics and solvent reorganisation. In cases where there are imperfections in the fiducial end-points, these are imprinted on the trajectory. Therefore, substantial care should be taken to check Ramachandran outliers and regions of poor geometry before applying this method. This helps to prevent inaccurate conclusions being drawn from poorly modelled protein structures. In cases where structural data is available [25], we show that is it possible to improve the model and rectify true outliers prior to calculating the trajectory as applied here for MATE. By cold-inbetweening the trajectory of transitions of protein structures, we can generate testable hypotheses as to how the scaffold of the protein allows these transitions to contribute to the protein’s function.

## Supporting information

trajectories

## 5 Acknowledgements

We thank David Stuart, Stephen McCarthy, Godfrey Boddard and Ilme Schlichting for comments on this paper and providing valuable feedback. H.M.G. is funded by the Helmholtz Association, grant VH-NG-19-02 (Helmholtz Young Investigator Group).

### 5.1 Data and code availability

The code for performing the cold-inbetweening is included in git commit 9da67bd of the RoPE software (https://github.com/helenginn/rope). Ensemble models of the three transitions are included in the Supporting Material.

## 6 Author Contributions

B.A.Y. data analysis, co-wrote the paper. H.M.G. research conception, wrote the cold-inbetweening code, data analysis, co-wrote the paper.

## 8 Methods

### 8.1 Construction of parameter set

The set of torsion angles refers to all torsion angles in a protein required to recalculate the protein’s conformation. Each torsion angle varies as a function of *p*, which varies from 0 to 1 along the trajectory. The starting and ending structures of the protein are fixed. At *p* = 0, each torsion angle *t*_*i*_ in the structure is defined to have the starting value 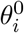, and at *p* = 1, the ending value 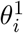. Each of the *t*_*i*_ values for *i* = 0 … *n* is a linear sum of terms as a function of *p*. The first of these terms (order 0) is a linear interpolation as in Eq. 1 and has no parameters.

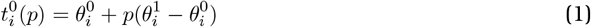

Additional terms are defined for the maximum order 3. *f*_*n*_ for each *n* is an amplitude and a refineable parameter, for which the default value is zero.

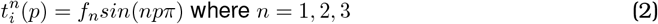

These are combined to produce an interpolation for each torsion angle with 3 refineable parameters,

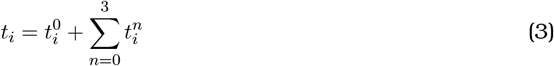

### 8.2 Torsion angle direction choice

For each torsion angle, the direction of clockwise or anticlockwise must be chosen, which is not necessarily evident from the start or end states. Torsion angles which differ by no more than 30*°* between start and end state are considered to move 30*°* rather than 330*°*. For all other torsion angles, an algorithm is employed to choose the more appropriate direction within the context of the rest of the molecule.

Each atom has a starting position, 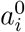, and an ending position 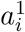. At a position *p* along the trajectory, an atom has a linearly interpolated position 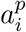:

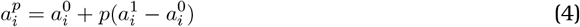

Sets of four atoms associated with each torsion angle were assigned consecutive values of 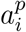 for values of *p* = 0, 0.1, 0.2……1.0. For the first iteration, a reference torsion angle was calculated from these sets of four atoms. For subsequent iterations, iteration *n*, an updated torsion angle was similarly calculated, and either -360, 0 or 360 was added in order to minimise the difference with the reference torsion angle from iteration *n* − 1. This torsion angle is then carried through for the next calculation for iteration *n* + 1. The final torsion angle for *p* = 1.0 is taken to indicate the direction of travel for this atom.

### 8.3 Construction of early-stage target function

Each atom has a starting position and ending position, which are a function of the set of torsion angles denoted by 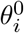 and 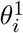 over all *i*. This means that every pair of non-hydrogen atoms *i* and *j* have a calculable distance 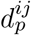 for a given value of *p*, corresponding to an invariant beginning and ending distance at *p* = 0 and *p* = 1 respectively. Every pair of non-hydrogen atoms for which either 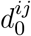 or 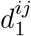 is below a fixed limit chosen to balance coverage of close contacts and computational efficiency (default value of 8 Å), and which are not covalently connected by three or fewer intervening bonds, is included in the early-stage target function.

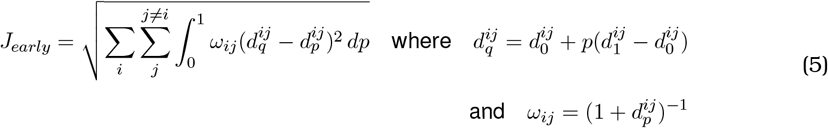

This therefore forms a least squares target function between the target atomic distances or glide-paths 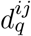, which are a linear interpolation with respect to *p*, and the calculated atomic distances 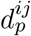 from the set of torsion angles for each value of *p*. In practice, *J*_*early*_ is numerically integrated with a number of steps which is set to 12 by default. The *ω*_*ij*_ term upweights close contacts and helps escape overlapping protein backbones.

### 8.4 Construction of late-stage target function

The late-stage target function measures Van der Waals contacts and torsion angle energies, calculated over the pairs of atoms (including hydrogens) similarly below a fixed limit, which may need to be larger than 8 Å depending on the scale of the conformational change.

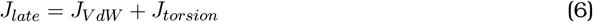

*J*_*V dW*_ consists of a discontinuous function measuring Van der Waals contacts above those expected from the target atomic distances using the two-term Lennard-Jones potential. This consists of a reference Van der Waals energy term *V dW*_*q*_(*ij*) for the interpolated distance, here *ϵ* is the potential well depth and a measure of the interaction strength [28] and *σ*^*i*^ is the van der Waals radius for atom *i* multiplied by 0.75.

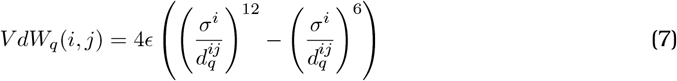

and a Van der Waals energy term *V dW*_*p*_(*ij*) for the calculated distance,

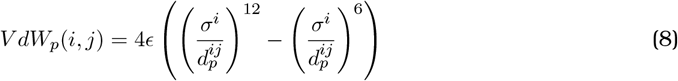

The *V dW*_*q*_ term is often positive for hydrogens involved in hydrogen-bonding arrangements which should not necessarily be disturbed during the transition. Therefore, it is used as a threshold, so only the positive difference between *V dW*_*p*_ and *V dW*_*q*_ is considered.

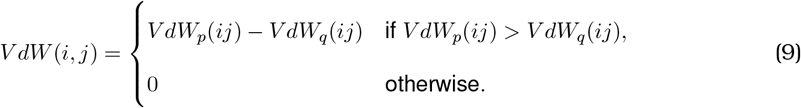

The Van der Waals component of the late-stage target function is calculated as follows.

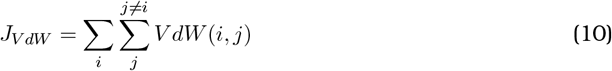

Consider the pair of atoms (A, B) for each bond in the structure. A torsion angle will connect four atoms in total. Each bond will be the centre of a number of related torsion angles. To generate a general torsion angle energy function which approximates the energies associated with eclipsed and staggered conformations, each torsion angle is considered in turn. We find the set of *n* torsion angles centred on each bond. We then define the total torsion energy as a sum of each part,

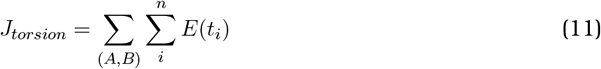

where *E*(*t*_*i*_) is the corresponding energy term of the *i*th torsion,

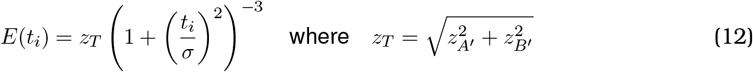

In this bell-curve function, *z*_*A*_*′* and *z*_*B*_*′* are the atomic numbers of the terminal atoms of the torsion angle directly bonded to A and B, respectively. *σ* denotes a bell curve width set to 60*°*.

*J*_*late*_ is numerically integrated with the same 12 steps for path calculations, but with some adjustments in *J*_*V dW*_. By nature of slicing of the pathway into intervals for numerical integration, sometimes atoms may pass extremely close to each other, but the point of minimum distance falls in between two numerically integrated intervals. This would create a way in which a minimisation method can easily mis-fit, in conjunction with a rapidly-changing function like *J*_*V dW*_. Therefore, a modification to this approach was taken: for an atom moving between intervals *n* and *n* + 1, *n* − 1 and *n* + 2 intervals were also used to generate a cubic spline in dimensions *x, y, z* where the polynomial must pass through each of these atom positions. When comparing the trajectories of atoms *i* and *j*, the minimum distance between these atoms between fractional values between *n* and *n* + 1 are found and this value is taken as the interval’s partial sum towards the estimation of the total integral.

### 8.5 Minimisation protocols

Minimisation against the *J*_*early*_ term is carried out using the L-BFGS algorithm [29]. Minimisation is considered converged if there is less than a 10^−3^ change in the *J*_*early*_ term after a L-BFGS refinement cycle. Gradient descent is carried out on *f*_1_ for the set of all torsion angles until convergence is reached; then on *f*_1_, *f*_2_ until convergence is reached, and likewise for parameters *f*_1_, *f*_2_, *f*_3_.

Minimisation against *J*_*late*_ is carried out using the Nelder-Mead simplex descent algorithm [30]. As each sidechain is almost independent from the rest of the protein, barring other local sidechains, simplex descents are run for each sidechain concurrently. Parameters *f*_1_, *f*_2_, *f*_3_ are refined for torsion angles which affect the sidechain and do not affect the main chain with a step size of 10*°*. After this, residues are ranked in terms of descending *J*_*late*_ terms, including all atoms contributing to that residue’s side or main chain. This is followed by additional simplex descents for the five residues with the highest values of *J*_*late*_ term. This two-stage process is repeated until the *J*_*late*_ term does not decrease by more than 5%.

### 8.6 Manual adjustments of torsion angles

Due to the subtlety of interactions between torsion angle directional choices, manual adjustment of a handful of parameters is required. Some backbone torsion angles must be manually adjustedwwhere many neighbouring bonds frequently exceed 90*°*, as this is easily confusable with a 270*°* rotation in the opposite direction. These decision-making processes were aided by a GUI which visually displays atom trajectories and makes bond flipping and harmonic adjustment accessible to users. Harmonic adjustment was occasionally employed during Nelder-Mead minimisation in order to escape Van der Waals “tangled” side chains, which are trapped in local, but not global, minima due to crossed bonds.

### 8.7 Preparation of structures

Structures were downloaded from PDB-REDO [31] to regularise the use of software used to refine structures. Structures were re-hydrogenated using *phenix*.*reduce*. These structures were prepared for cold-inbetweening using the RoPE software package [32] which optimises torsion angles to fit the experimentally determined model [12]. Before cold-inbetweening, to ensure the simplest transition, the torsion angles of 180*°*-symmetric side chains (phenylalanine, tyrosine, aspartate, glutamate, arginine) were made identical. Due to the lower resolution (*>* 2.4 Å) of structures involved, this was also extended to residues which could be distinguished only at higher resolution (asparagine, glutamine, histidine). Cold-inbetweening is implemented in the RoPE software package.

### 8.8 Cavity calculations

The calculation of the ligand cavity for EIIC domain of the MalT transporter was performed using the KVFinder web server [27]. The molecular surface of the protein was calculated using a 1.7 Å probe radius and the inaccessible regions were calculated using a 4 Å probe radius. The outer limits of the cavity were defined using a 2.4 Å removal distance. The cavity volume cutoff was set according to the molecular volume of maltose 241.54 Å^3^. The molecular volume of maltose was calculated with MoloVol [33], using a carbon atom radius of 1.77 Å and oxygen radius of 1.50 Å, with a probe of 1.2 Å, grid resolution 0.2 Å and optimisation depth 4.

**Supporting Fig. 1:**
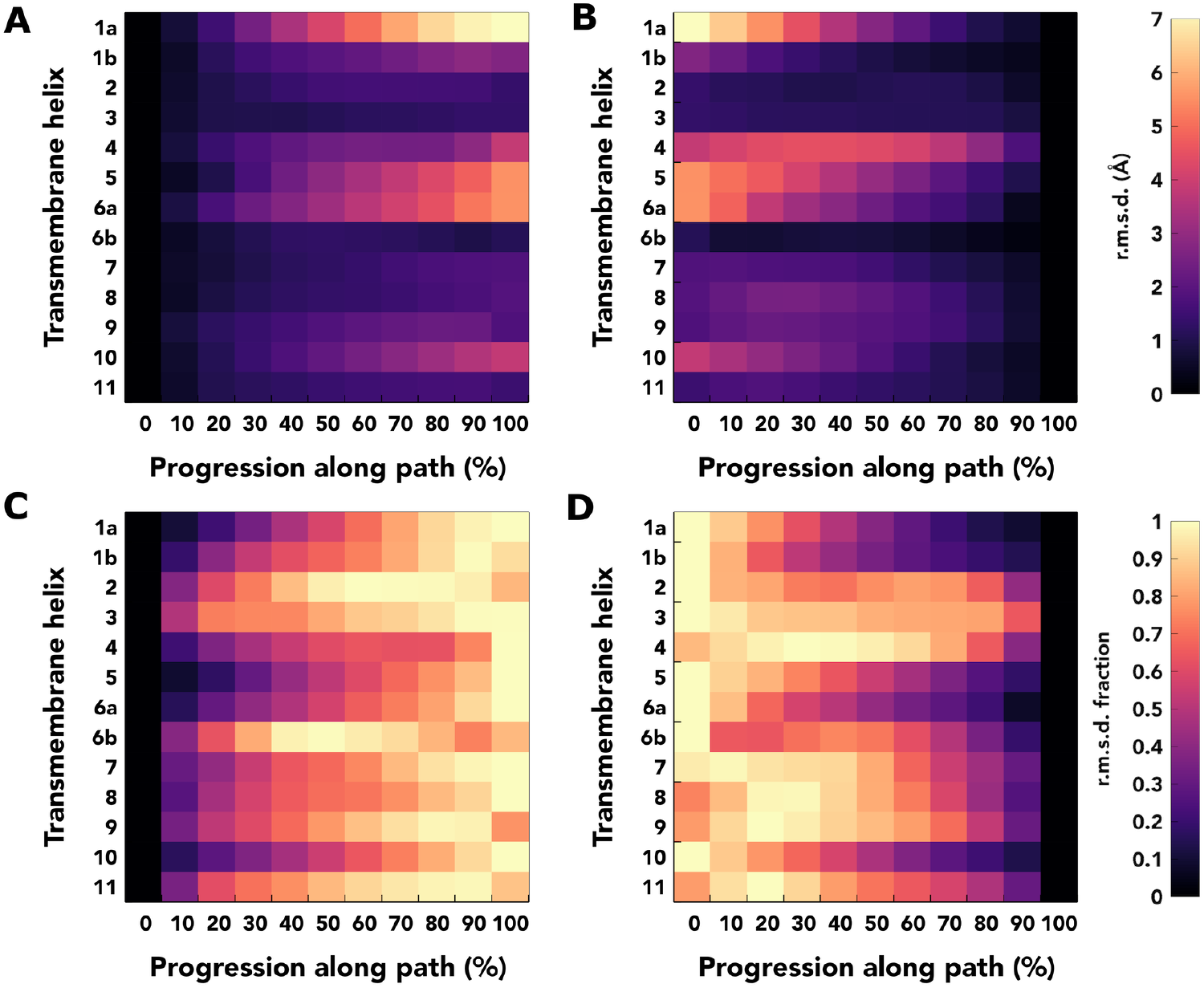
Helix alignments of mid-trajectory points along the pathway with the start and end points of the DraNramp cold-inbetweened pathway. (A, B) alignment in absolute r.m.s.d. against (A) 0% of trajectory and (B) 100% of trajectory. (C, D) alignment where each helix alignment is expressed as a fraction of the maximum r.m.s.d. for that helix, against (C) 0% trajectory and (D) 100% trajectory, respectively.

**Supporting Fig. 2:**
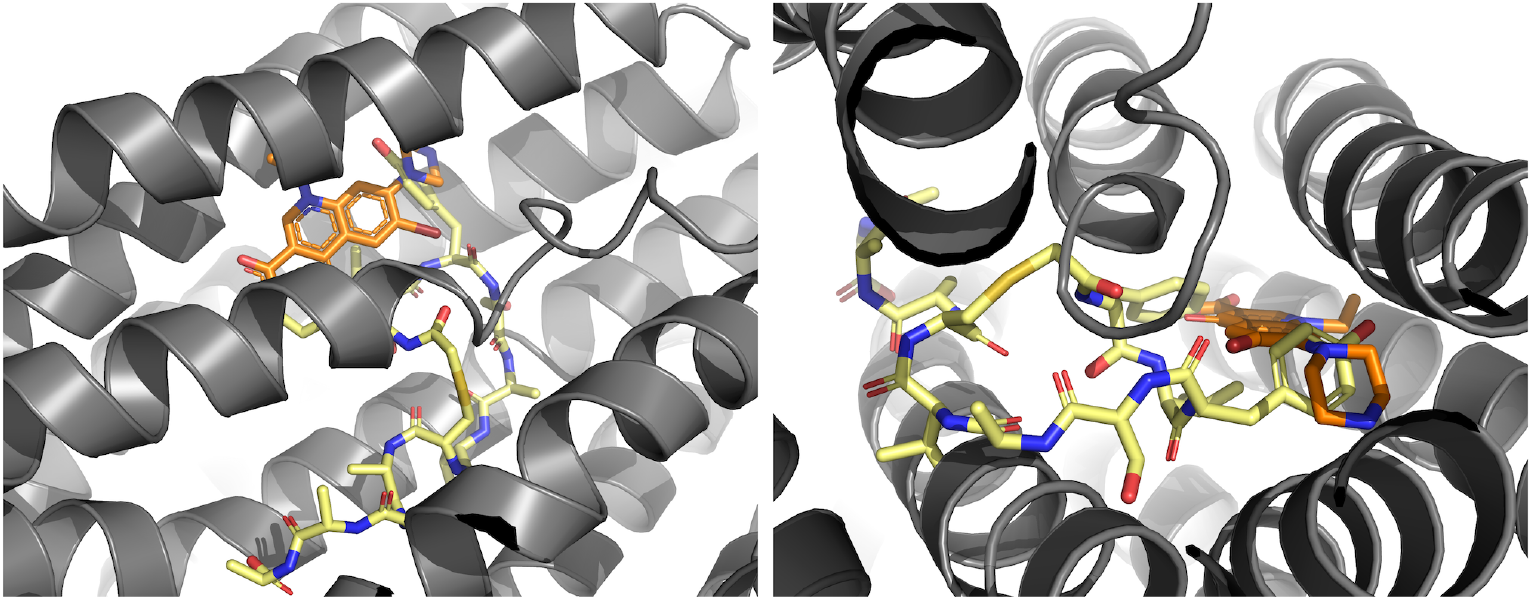
Two views of 70% progression (gray) through the MATE transition from inward-to outward-open, at the kinked TM1 unwinding position. Overlaid are the Br-NRF ligand-bound structure (orange) and the MaD5 inhibitor-bound structure (yellow) using all atoms in residues 165-185 for structure alignment (TM3, adjacent to ligand-binding site).

